# Opposing associations of Internet Use Disorder symptom domains with structural and functional organization of the striatum: a dimensional neuroimaging approach

**DOI:** 10.1101/2022.05.07.491044

**Authors:** Fangwen Yu, Jialin Li, Lei Xu, Xiaoxiao Zheng, Meina Fu, Keshuang Li, Shuxia Yao, Keith M Kendrick, Christian Montag, Benjamin Becker

## Abstract

**Background:** Accumulating evidence suggests brain structural and functional alterations in Internet Use Disorder (IUD). However, conclusions are strongly limited due to the retrospective case-control design of the studies, small samples, and the focus on general rather than symptom-specific approaches.

**Methods:** We here employed a dimensional multi-methodical MRI-neuroimaging design in a final sample of n = 203 subjects to examine associations between levels of IUD and its symptom-dimensions (loss of control/time management, craving/social problems) with brain structure, resting state and task-based (pain empathy, affective go/no-go) brain function.

**Results:** Although the present sample covered the entire range of IUD, including normal, problematic as well as pathological levels, general IUD symptom load was not associated with brain structural or functional alterations. However, the symptom-dimensions exhibited opposing associations with the intrinsic and structural organization of the brain, such that loss of control/time management exhibited negative associations with intrinsic striatal networks and hippocampal volume, while craving/social problems exhibited a positive association with intrinsic striatal networks and caudate volume.

**Conclusions:** Our findings provided the first evidence for IUD symptom-domain specific associations with progressive alterations in the intrinsic structural and functional organization of the brain, particularly of striatal systems involved in reward, habitual and cognitive control processes.

## Introduction

Internet use disorders (IUD) have become a growing public health concern. IUD is characterized by excessive Internet usage, loss of control over usage and detrimental consequences for occupational and social functioning. IUD represents an umbrella term encompassing Internet-associated use disorders, i.e. Internet gaming disorder (IGD) or Social Networks Use Disorder (Montag et al., 2021a; b; Montag, Schivinski & Pontes, 2021; Musetti et al., 2016; Spada, 2014). Estimated prevalence rates of IUD or its subforms such as IGD range for from 1% to 10%, with particularly high rates among young individuals and in Asia with up to 25% meeting the criteria for some forms of IUD (Pan, Chiu, & Lin, 2020; for Asia see Montag & Becker; 2020). Debates continue on whether IUD represents a mental disorder (Musetti et al., 2016; Starcevic & Khazaal, 2017), yet (I)GD has been included in the current diagnostic classification systems (DSM-V, ICD-11) and conceptual perspectives propose symptomatic and neurobiological similarities with behavioral and substance use addictions (Montag & Reuter, 2017; Musetti et al., 2016; Young, 1998b, 2011).

Several psychometric instruments have been developed to assess specific forms of IUD, including IGD or the use of specific platforms (Montag et al., 2018; Pontes et al., 2021), while the extensively validated Internet Addiction Diagnostic Questionnaire and the Internet addiction test (IAT) measure level of IUD in general (Young, 1998a, 1999). The symptoms assessed by these scales strongly resemble symptoms of substance addiction, such that the items of the IAT assess preoccupation with the Internet, withdrawal, loss of regulatory control and social problems (Musetti et al., 2016; Young, 1998b). While different factorial structures of the IAT ranging from one to six factors have been proposed based on exploratory factor analyses (Chang & Law, 2008; Khazaal et al., 2008; Korkeila, Kaarlas, Jääskeläinen, Vahlberg, & Taiminen, 2010; Widyanto, Griffiths, & Brunsden, 2011), confirmatory factor analyses consistently revealed a two-factorial model of the IAT across Western and Chinese samples (Pawlikowski, Altstotter-Gleich, & Brand, 2013; Stodt et al., 2018). The factors describe the key facets of ‘loss of control/time management’, which strongly predicts obsessive-compulsive and impulsivity symptoms and ‘craving/social problems’, which predicts emotional and interpersonal dysregulations. The two factorial solution has been further confirmed by good convergent, divergent and incremental validity of the two-factor structure of the short version of the IAT (s-IAT) across independent samples (Pawlikowski et al., 2013; Stodt et al., 2018).

In line with the similarities on the symptomatic level, accumulating evidence suggests that functional and structural brain changes associated with problematic internet use partly resemble alterations observed in substance addiction (Klugah-Brown et al., 2020; Klugah-Brown et al., 2021; Tolomeo & Yu, 2022). Similar to substance use disorders, IGD showed reduced gray matter volumes in the striatal reward system, as well as insular, frontal and temporal systems involved in executive control, decision-making and social processes (He, Turel, & Bechara, 2017; Montag et al., 2018; Zhou et al., 2011; Zhou et al., 2019a). Functional MRI studies reported cue and craving associated hyperreactivity in striatal, cingulate and lateral prefrontal cortex (DLPFC) regions (Han, Hwang, & Renshaw, 2011; Ko et al., 2009; Ko et al., 2013a; Ko et al., 2013b; Lorenz et al., 2013) while cognitive and emotional control impairments were associated with abnormal engagement of fronto-parietal, insular and temporal regions (Chen et al., 2015; Ding et al., 2014; Dong, Devito, Du, & Cui, 2012; Dong, Zhou, & Zhao, 2010; Ko et al., 2014; Liu et al., 2014; Shin, Kim, Kim, & Kim, 2021; Turel, He, Xue, Xiao, & Bechara, 2014; Zhang et al., 2020). Alterations in the intrinsic functional organization of the brain have been extensively examined in individuals with problematic Internet use by means of resting-state fMRI with recent large scale and meta-analysis studies reporting altered functional connectivity in striato-insular-frontal circuits (Dong et al., 2021a; Yan, Li, Yu, & Zhao, 2021).

However, the findings for IUD (particularly for IGD) have been inconsistent, such that prospective longitudinal studies could not confirm emotional or empathic alterations previously observed in cross-section designs (Anderson et al., 2008; Anderson et al., 2010; Gao et al., 2017; Jiao, Wang, Peng, & Cui, 2017; Kühn et al., 2018). These inconsistencies may partly be related to the retrospective categorical case-control design of the studies that commonly compared relatively small sample sizes of individuals with excessive internet engagement or established IUD with healthy controls. The assumption of the case-control approach is that categorical diagnosis, e.g. fulfilling at least five of the DSM-5 IGD criteria or an IAT score >50 will define mechanistically meaningful study samples, whereas in fact the categorical IUD groups neither exhibit the same symptom constellation nor cover the whole symptom spectrum of IUD (Etkin, 2019). Several of the early studies employ sample sizes between 20 to 30, or often less than 20 per group (Cheng & Liu, 2020; Dong et al., 2010; Lorenz et al., 2013), which permits statistical detection power to be achieved but at the cost of sacrificing clinical significance.

Recent suggestions to improve psychiatric neuroimaging advocate prospective dimensional approaches that examine associations between varying degrees in symptom load and progressive neural alterations in larger samples (Etkin, 2019). An increasing number of studies successfully employed this approach to determine symptom-dimension specific brain alterations e.g. with respect to depression or alexithymia (Hägele et al., 2016; Li et al., 2019; Luo et al., 2018) and initial studies have demonstrated the feasibility of this approach to determine IUD-associated brain structural and intrinsic functional changes in larger samples spanning the entire symptom load range (Dong et al., 2021a; Montag et al., 2018; Zhou et al., 2021). Against this background we employed a dimensional multi-modal neuroimaging approach in a comparably large sample (> 240 subjects) to explore associations between the level of problematic Internet use (as assessed by the IAT) on brain structure as well as intrinsic and task-associated brain function. We focused on task domains for which IUD alterations are a matter of debate, in particular cognitive control and implicit emotion regulation (affective go/no-go) and empathy or social communicative processing (pain empathy task including physical and social communicative pain). To further disentangle the impact of different symptom constellations which cannot be examined with the categorical approach the symptom dimensions “loss of control/time” and “craving/social problem” (Pawlikowski et al., 2013) were separately examined (Stodt et al., 2018). In line with previous studies, we expected that higher levels of problematic Internet use would be associated with volume and intrinsic striatal connectivity changes as well as deficient frontal and insular engagement during cognitive control and empathic processing, respectively. With respect to the sub-facets we expected distinguishable symptom-specific associations.

## Methods

### Participants

To implement dimensional neuroimaging in a large sample we capitalized on the Chengdu Gene Brain Behavior Project which aims at determining psychopathological, genetic and neural alterations in a large cohort of healthy individuals (Liu et al., 2021; Montag et al., 2017b). N = 250 subjects from this project underwent multimodal MRI imaging, including a pain empathy paradigm (Li et al., 2019; Xu et al., 2020; Zhou et al., 2020a), an affective Go/No-go paradigm (Zhuang et al., 2021) as well as resting state and brain structural assessments (e.g. Liu et al., 2021). Levels of IUD were assessed at the time of the MRI-data acquisition by means of the Internet Addiction Test (IAT, Pawlikowski et al., 2013; Stodt et al., 2018; Young, 1998b) which assess a total score as well as the sub-facets describing loss of control/time management and craving/social problems in the context of problematic Internet use.

Following data quality assessments N=203 subjects were included in the final multimodal data analyses (104 males; mean age ±SD=21.65 ±2.37 years). Participants provided written informed consent, the study had full ethical approval and was in accordance with the latest revision of the declaration of Helsinki.

### Experimental Protocols

Internet addiction and the sub-facets were assessed using the short version of Young’s Internet Addiction Test (s-IAT, Pawlikowski et al., 2013). The s-IAT includes 12-items which assess the frequency of negative experiences and consequences due to excessive online activities in everyday life on a 5-point Likert scale ranging from 1 (never) to 5 (very often) (Young, 1998). The total score can correspondingly range from 12 to 60, with a total score >30 indicating problematic and >37 indicating pathological Internet use (Pawlikowski et al., 2013; Stodt et al., 2018). The s-IAT consists of the two subscales assessing loss of control/time management and craving/social problems each consisting of six items (items see table S1). Subjects underwent MRI assessments, including acquisition of T1-weigted brain structural data, resting state fMRI data as well as a validated fMRI pain empathy task and an affective Go/No-go task (details see supplemental material and figure 1).

**Figure 1.**
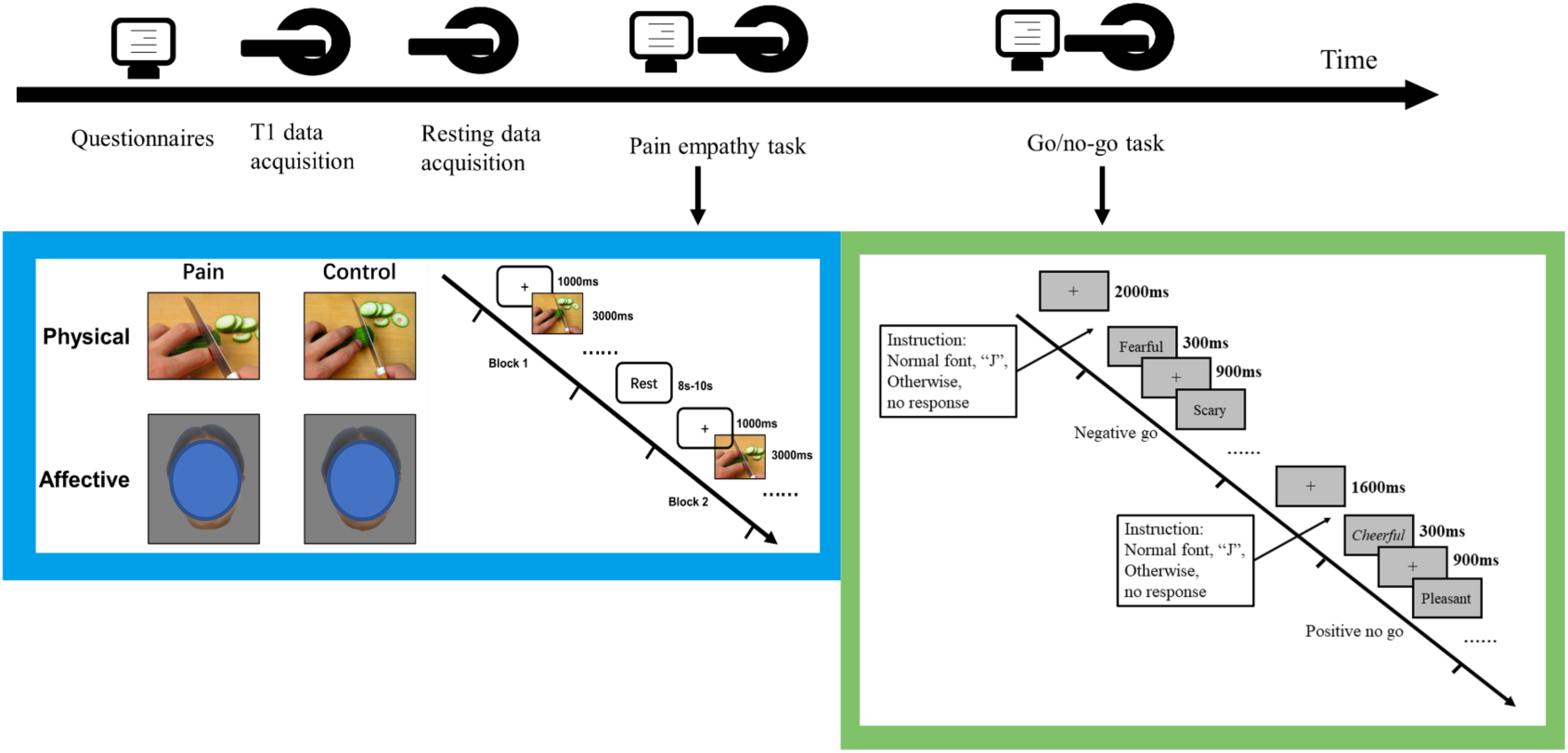
Experimental procedure (note: for the preprint version the faces have been overlayed with a blue circle)

### MRI Data Processing, Dimensional Approach and Thresholding

MRI data were acquired and processed using extensively validated standard procedures and processing pipelines (see supplements). We employed a dimensional neuroimaging approach modelling associations between the severity of Internet addiction (aka Internet Use Disorders) and its sub-facets (as assessed by the s-IAT) and individual variations in brain function and structure using voxel-level multiple regression analyses in SPM12. To this end the s-IAT total and subscale scores served as predictor while the voxel-wise brain structural, task activation and intrinsic connectivity maps served as dependent variable. Given that the s-IAT total scores consisted of the sum of loss of control/time management subscale scores and craving/social problems subscales separate multiple regression analyses for the total scores and two subscales, both including age and sex as covariates of no interest (in the VBM analysis also TIV) were conducted. Analyses were conducted on the whole-brain level with FDR *p* < 0.05 for multiple comparison correction. Details on the paradigms, preprocessing and first level models are provided in the supplements. The voxel-wise whole-brain maps were modelled as follows.

### Task fMRI pain empathy

The paradigm included visual stimuli depicting ‘physical pain’, ‘affective pain’ and corresponding non-painful control stimuli (‘physical control’ and ‘affective control’). The first level matrix included separate regressors for the four experimental conditions and six head-motion parameters. In the absence of empathy type-specific hypotheses we focused on the main pain empathy contrast [(physical pain + affective pain) > (physical control + affective control)], with further exploratory analyses examining associations with empathy-type specific activity ([physical pain > physical control]; [affective pain > affective control]).

### Task fMRI affective Go/No-go

The lexical Go/No-go paradigm required go and no-go responses in neutral, negative and positive contexts leading to six experimental conditions on the first level (neu Go, neu No Go, neg Go, neg No Go, pos Go, pos No Go) and additional six head-motion parameters. We focused on the general inhibitory control contrast [(neu No Go + neg No Go + pos No Go) > (neu Go + neg Go + pos Go)], with further exploratory analyses exploring associations with emotional context-specific inhibitory activity ([neu No Go > neu Go], [neg No Go > neg Go], ([pos No Go > pos Go]). During the paradigm accuracy and reaction times were collected and served as behavioral outcomes.

### Resting state functional connectivity

Based on the different roles of the ventral and dorsal striatum in (internet) addiction, (Dong et al., 2021a; Dong et al., 2021b; Zhou et al., 2018; Zhou et al., 2020b; Zhou et al., 2019b) bilateral ventral striatum and dorsal striatum masks served as seed regions (regions of interest, ROIs). ROIs were derived from the brainnetome atlas (Fan et al., 2016) (table S4) functional connectivity between the ROIs and voxels on the whole brain level were computed using DPABI (Yan, Wang, Zuo, & Zang, 2016; https://www.rfmri.org/dpabi).

### Brain structure gray matter volume

Voxel-based morphometry (VBM) analysis was conducted using the CAT toolbox (version 12.7, http://www.neuro.uni-jena.de/cat/) including quality assessments, smoothing and statistical analysis. T1 images were normalized to a template space and segmented into gray matter (GM), white matter (WM), and cerebrospinal fluid (CSF). The preprocessing parameters used the default recommendations (“Segment Data”). After the preprocessing, we conducted a quality check and deleted data with mean correlation of sample homogeneity below two standard deviations. After segmentation, images were smoothed with 8mm FWHM and resulting voxel-wise gar matter masks were subjected to the multiple regression models.

## Results

### Participants and levels of IUD

Mean total score of the s-IAT was 32.49 (SD=7.62) in the present sample (N=203). 74 subjects scored below 30 indicating normal Internet use, n=81 subjects reported scores between 30 and 37 suggesting problematic Internet use, while 48 subjects reported total scores >37 suggesting pathological Internet use (Pawlikowski et al., 2013; Stodt et al., 2018). With respect to sex differences, the total levels, *t*=1.06, *df*=201, *p*=0.289, Cohen’s *d*=0.149, and levels of craving/social problem were comparable *t*=0.757, *df*=201, *p*=0.450, Cohen’s *d*=0.106 while for the loss of control/time management females reported higher levels than males (*t*=2.682, *df*=201, *p*=0.008, Cohen’s *d*=0.377).

Sex was consequently included in all analyses as covariate. Details see table 1. The data displayed no strong violations of normal distribution regarding the s-IAT and its subscales. Moreover, the scales showed good psychometric properties (details see table S2, supplements, supplementary figure S1)

**Table 1.** General and symptom-domain levels of Internet Use disorders as measured by the s-IAT and its subscales

| Questionnaire | Gender | Mean (SD) | *t* | *p* | Cohen's *d* |
| --- | --- | --- | --- | --- | --- |
| Total scores | female | 33.07(8.01) | 1.06 | 0.289 | 0.149 |
|  | male | 31.93(7.23) |  |  |  |
| Loss of control subscale scores | female | 19.12(4.62) | 2.68 | 0.008 | 0.377 |
|  | male | 17.55(3.71) |  |  |  |
| Craving/social problem subscale scores | female | 13.95(4.03) | 0.44 | 0.450 | 0.106 |
|  | male | 14.38(4.15) |  |  |  |

### Associations between levels of IUD and affective Go/No-Go performance

Partial correlation analyses with accuracy and reaction time of go and no-go trials for the three conditions (negative, positive, neutral) and age and gender as covariates revealed that the s-IAT total scores and two subscale scores had significant negative correlations with the accuracy in no-go trials (Negative no-go: *r*_total_=-0.215, *p*=0.00215, *r*_control_=-0.186, *p*=0.00823, *r*_craving_=-0.211, *p*=0.00266; Positive no-go: *r*_total_=-0.233, *p*=0.00087, *r*_control_=-0.189, *p*=0.00717, *r*_craving_=-0.241, *p*=0.00058; Neutral no-go: *r*_total_=-0.203, *p*=0.00391, *r*_control_=-0.162, *p*=0.02137, *r*_craving_=-0.211, *p*=0.00258, Bonferroni corrected *p*=0.0083, six tests). No significant correlation with accuracy and reaction times of go trials were observed after Bonferroni correction. Details see table S3.

### Associations between levels of IUD and pain empathy and cognitive/emotional control associated brain activity

The pain empathy paradigm engaged per se the typical bilateral pain empathy networks encompassing e.g. the bilateral inferior frontal gyrus (IFG) and insula and medial frontal cortex (FDR *p*<0.05, cluster size>297, see table S5 and figure S2A). The go/no-go paradigm engaged the key frontoparietal inhibitory control networks (contrast [all no/go > all go], FDR *p*<0.05) (figure S2B). For details on other contrasts please see also Zhou et al., 2020(a); Zhuang et al., 2021. Multiple regression analysis examining associations with IUD total and subscale scores did not reveal significant associations with pain empathy and general inhibitory control associated brain activity (FDR *p*<0.05). Further exploratory analyses did not reveal significant associations with empathy-type or emotional specific inhibitory activity.

### Associations between levels of IUD and intrinsic striatal networks

While no associations were found for the total IAT score, the subfacets showed associations with striatal intrinsic connectivity networks. Levels of loss of control/time management were significantly negatively associated with the intrinsic functional connectivity of ventral striatum with the right calcarine, right middle occipital lobe, left superior occipital lobe and right fusiform gyrus (see table 2 and figure 2A, FDR *p*<0.05, cluster size>=49 voxels), while they were significantly negatively associated with dorsal striatum intrinsic connectivity with the right precuneus, right posterior cerebellum lobe and anterior cingulate cortex (see table 2 and figure 2A, FDR *p*<0.05, cluster size>=36 voxels). Levels of craving/social problems were significantly positively associated with functional connectivity of the ventral striatum with the right fusiform gyrus, left superior occipital lobe, right middle occipital lobe, left inferior orbital frontal cortex, right superior occipital lobe, right superior temporal lobe and left insula (see table 3 and figure 2B, FDR *p*<0.05, cluster size>=22 voxels). With respect to the dorsal striatum connectivity levels of craving/social problems were positively associated with coupling with the left occipital lobe, left inferior frontal gyrus and right inferior frontal gyrus (see table 3 and figure 2B, FDR *p*<0.05, cluster size>=27 voxels).

**Table 2.**
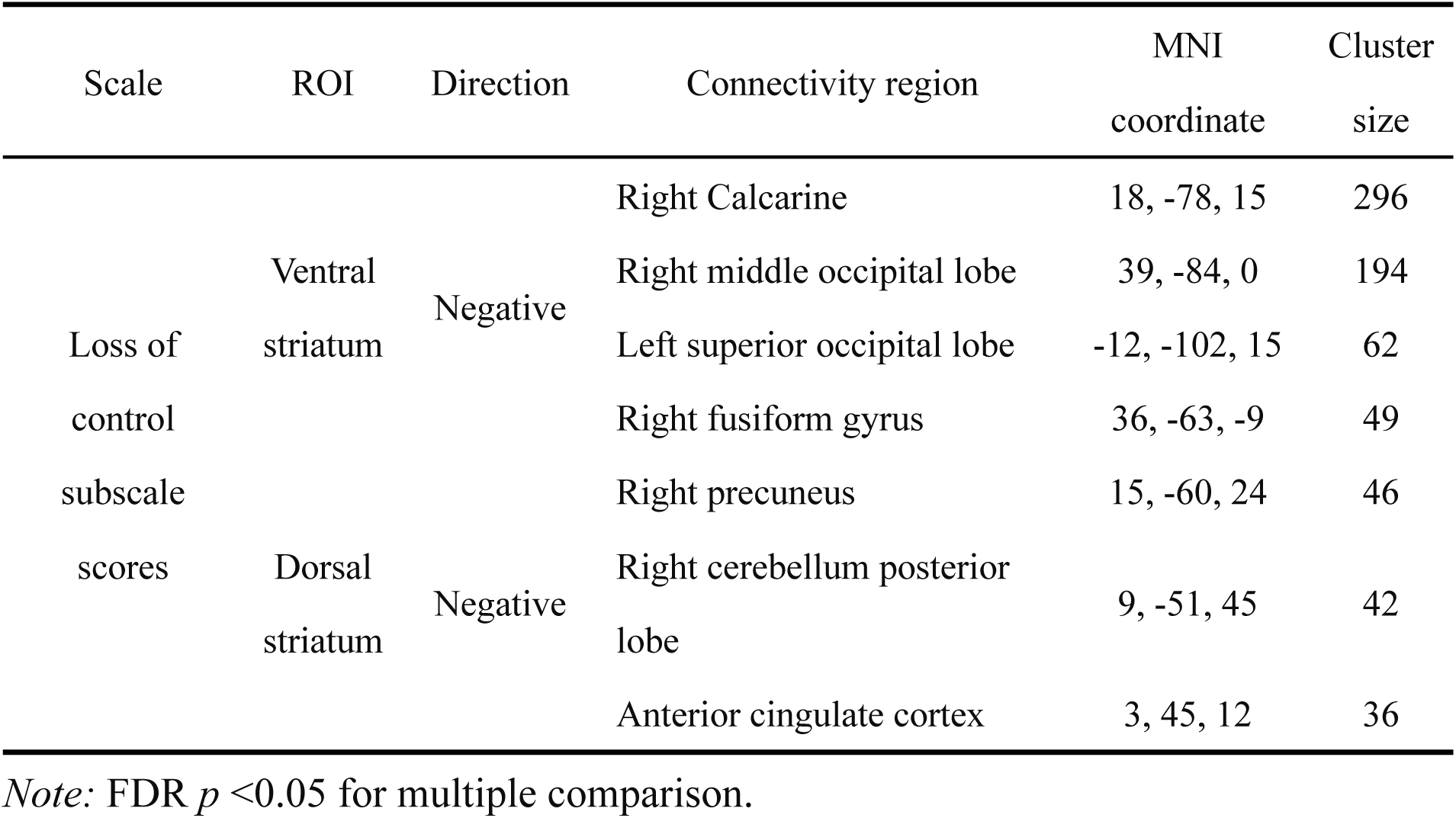
Associations between loss of control/time management and the striatum resting state networks

| Scale | ROI | Direction | Connectivity region | MNI coordinate | Cluster size |
| --- | --- | --- | --- | --- | --- |
| Loss of control subscale scores | Ventral striatum | Negative | Right Calcarine | 18, -78, 15 | 296 |
|  |  |  | Right middle occipital lobe | 39, -84, 0 | 194 |
|  |  |  | Left superior occipital lobe | -12, -102, 15 | 62 |
|  |  |  | Right fusiform gyrus | 36, -63, -9 | 49 |
|  | Dorsal striatum | Negative | Right precuneus | 15, -60, 24 | 46 |
|  |  |  | Right cerebellum posterior lobe | 9, -51, 45 | 42 |
|  |  |  | Anterior cingulate cortex | 3, 45, 12 | 36 |

*Note:* FDR $p < 0.05$ for multiple comparison.

**Table 3.**
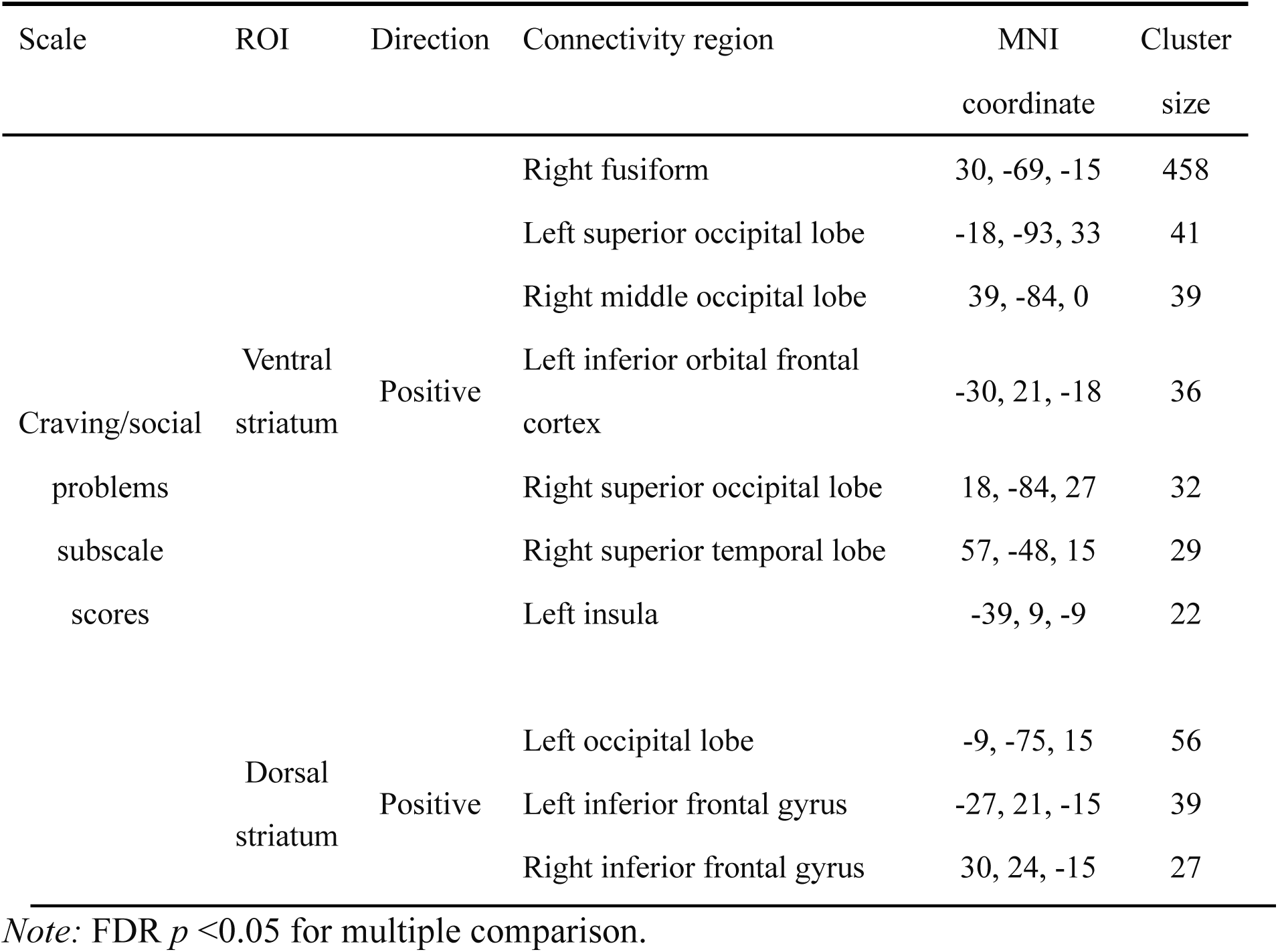
Associations between craving / social problems and striatal networks

**Figure 2.**
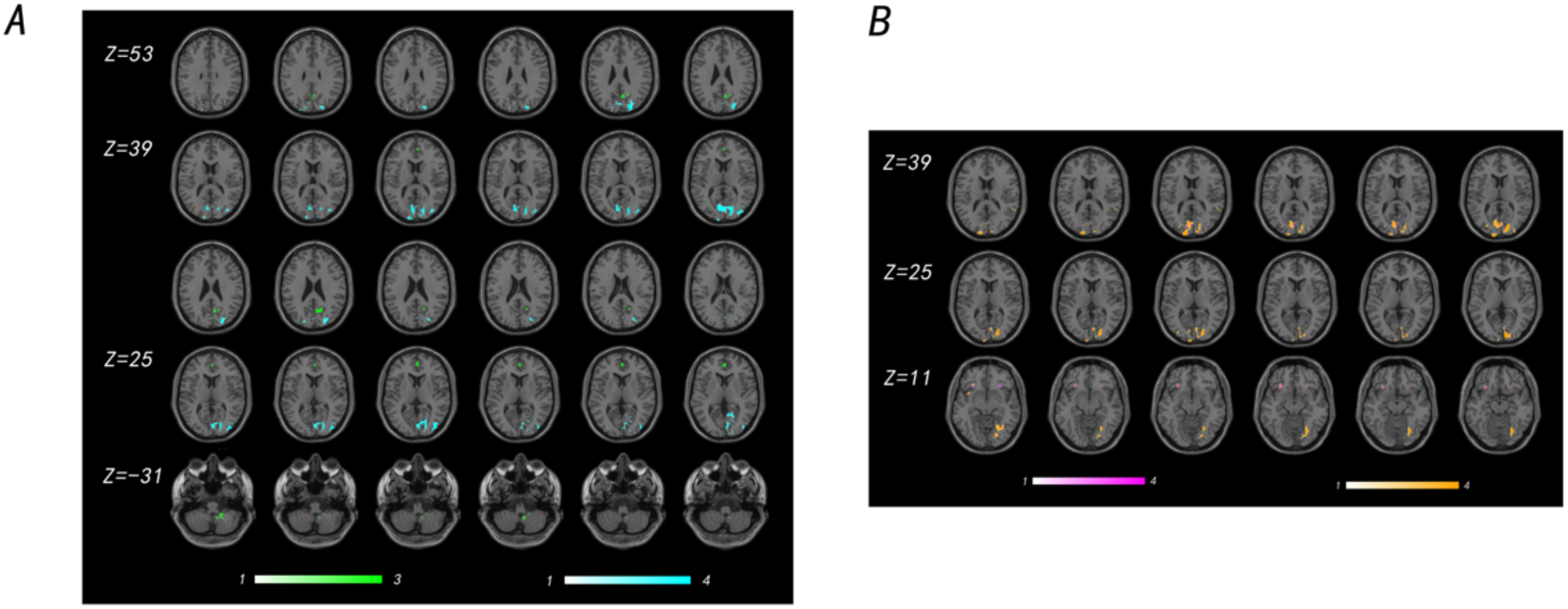
Associations of two subscale scores and striatum based resting state functional connectivity We firstly calculated the resting functional connectivity of bilateral ventral and dorsal striatum with voxels on the whole brain level, and conducted multiple regression analysis of connectivity, including loss of control subscale scores and craving/social problem subscale scores as covariates of interest, including age and gender as covariates of no interest. (A) The loss of control/time management subscale showed significant negative associations with resting state functional connectivity of the ventral striatum with right calcarine and right middle occipital lobe, left superior occipital lobe and right fusiform gyrus (FDR < 0.05, cluster size >= 49 voxels) (depicted in cyan). Loss of control/time management was significantly negative related to the resting state functional connectivity of dorsal striatum with right precuneus, right posterior cerebellum lobe and anterior cingulate cortex (FDR < 0.05, cluster size >= 36 voxels) (depicted in green). B: The craving/ significant positive association with resting state functional connectivity of the ventral striatum with right fusiform, left superior occipital lobe, right middle occipital lobe, left inferior orbital frontal cortex, right superior occipital lobe, right superior temporal lobe and left insula (FDR < 0.05, cluster size >= 22 voxels) (depicted in yellow). The craving/social problems subscale showed a significant positive relation with functional connectivity strengths of the dorsal striatum with the left occipital lobe, left inferior frontal gyrus and right inferior frontal gyrus (FDR < 0.05, cluster size >= 27 voxels) (depicted in magenta).

### Associations between levels of IUD and brain structure

No associations with s-IAT total score were observed but loss of control/time management exhibited a significant positive association with the GMV of the left hippocampus (MNI: -15/-18/-15, FDR *p*<0.05, cluster size=530 voxels, see figure 3A) while levels of craving/social problems showed a significant positive association with the GMV of the striatum (caudate subregion; MNI: 5/8/5, FDR *p*<0.05, cluster size=985 voxels, see figure 3B,). Mapping the caudate cluster using the brainnetome atlas revealed that the cluster encompassed ventral and dorsal striatal subregions (see table S6 and figure 3B).

**Figure 3.**
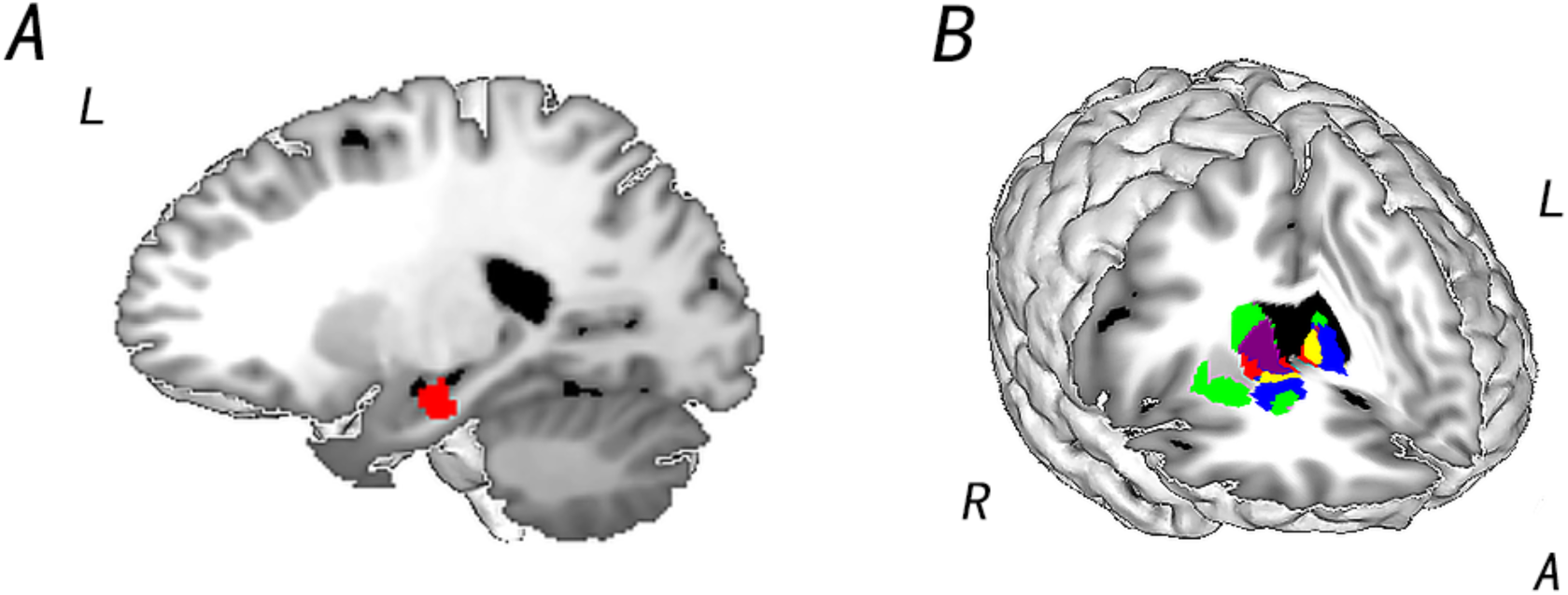
Associations of two subscales with gray matter volume on the whole brain level. A: Loss of control/time management was significantly positive associated with gray matter volume (GMV) of the left hippocampus (MNI coordinate: -15, -18, -15, FDR<0.05, cluster size = 530 voxels, see red region). B: Craving/social problems showed a significant positive association with GMV of Caudate (MNI coordinate: 5, 8, 5, FDR *p* <0.05, cluster size = 985 voxels, see red region). Purple represents the overlap with the caudate and bilateral dorsal striatum (green region); yellow represents regions representing the overlap between caudate and bilateral ventral striatum (blue region). Details see also table S7. *Note:* L: left; R: right; A: anterior.

## Discussion

We employed a dimensional neuroimaging approach in a comparably large sample and across different imaging modalities to determine early brain markers for IUD. Although a sample of young students was enrolled our sample spanned the entire range of IUD, with 81 or 43 from 203 individuals reporting problematic (n = 81) or pathological levels of internet use (n = 43), respectively (according to cut off scores provided in Pawlikowski et al., 2013; Stodt et al., 2018). Levels of IUD had negative associations with response inhibition performance in an affective go/no-go task in the absence of significant associations with brain activity during cognitive/emotional control or pain empathic processing. Although general severity of IUD (s-IAT total scores) did not associate with brain structural or functional organization, the sub-facets exhibited significant and opposing associations, such that loss of control/time management exhibited negative associations with intrinsic striatal networks and hippocampal volume, while craving/social problems demonstrated positive associations with striatal networks and caudate volume.

The dimensional approach allowed us for the first time to disentangle symptom-domain specific alterations, thus providing a neurobiological basis of previously reported differential associations with pathological domains (Etkin 2019; Pawlikowski et al., 2013) and suggesting that the behavioral dysregulations are mediated by separable neurobiological alterations. Higher levels of loss of control/time management symptom load were associated with decreasing connectivity of the ventral striatum with the occipital and fusiform regions and of the dorsal striatum with the precuneus and the anterior cingulate cortex. In contrast, higher levels of craving/social problems were associated with enhanced connectivity between the ventral striatum and the occipital, orbitofrontal and insular regions. Negative associations between loss of control and striatum connectivity might reflect dysregulations in inhibitory control and visual salience processing including the superior and middle occipital regions, and the action control system, including the cerebellum and ACC (Dong et al., 2021b; Meng, Deng, Wang, Guo, & Li, 2015). Positive associations between craving/social problems and the ventral striatal system encompassed occipital regions involved in visual salience processing and orbitofrontal regions involved in reward processing which have been associated with cue-reactivity and craving (Hanlon, Dowdle, Naselaris, Canterberry, & Cortese, 2014; Zhou et al., 2019b). The identified circuits moreover overlap with circuits reported to be involved in cognitive control deficits, reward dysregulations and cue-reactivity in both IUD (Ma et al., 2019; Starcke, Antons, Trotzke, & Brand, 2018; Yu et al., 2021; Yuan et al., 2017) and substance-use disorders (Montag & Reuter, 2017; Verdejo-Garcia, Garcia-Fernandez, & Dom, 2019; Volkow & Boyle, 2018). In substance addiction progressive adaptations in these circuits have been explained in terms of ‘incentive sensitization’ reflecting that the incentive motivational effects of drug and drug-associated stimuli leads to progressive dysfunctions in salience and reward processing and impaired executive control (Robinson & Berridge, 1993, 2008) or in terms of dysregulated ‘habit formation’, a process during which initially reinforced goal-directed actions become progressively habitual and compulsive (Ersche et al., 2016; Robbins & Clark, 2015). The excessive engagement in Internet usage may additionally be promoted by negative reinforcement learning such that these activities can reduce stress induced by societal and social pressure and attenuate anxiety and depression (King & Delfabbro, 2014, 2018).

The different neurobiological underpinnings of the two symptom scales were additionally mirrored on the brain structural level such that craving/social problems positively associated with the GMV of the striatum, including both ventral and dorsal parts, while higher levels of loss of control/time management problems were associated with larger GMV of the hippocampus. Previous studies reported increased GMV of the dorsal and ventral striatum in IUD and the volumes correlated with cognitive control performance and IAT scores (Cai et al., 2016). Although previous dimensional studies reported negative associations between problematic internet usage and striatal volume (Montag et al., 2017a; Montag et al., 2018; Zhou et al., 2020), morphological alterations in the mesocorticolimbic reward systems – also positive associations - and their structural connectivity have been extensively documented in IUD, in particular IGD (He, Turel & Bechara 2017; Hong et al., 2015; Wang et al., 2019; Zhou et al., 2019a; Zhou et al., 2011). In the context of the previous literature the current findings indicate that alterations in these pathways may specifically promote craving/social problems in IUD. In contrast, higher levels of control/time management were associated with larger hippocampal volumes. Previous studies have reported decreased as well as increased hippocampal volumes in IGD (Lin, Dong, Wang, & Du, 2015; Yoon et al., 2017) and this region has been suggested to play a role in the interaction between reward processing, behavioral reinforcement and memory retrieval (Everitt & Robbins, 2005).

This role in the development of IUD has been confirmed by a prospective longitudinal study employing gaming training (“Super Mario 64 DS”) at least 30 minutes per day over two months (Gleich, Lorenz, Gallinat, & Kuhn, 2017). Before and after training phase, both groups were shown gaming videos, which recorded during training, with three consequences (reward, punishment and neural) in the fMRI scanner. The results showed that gaming increased brain activity in the left hippocampus after training that reward processing during gaming may lead to a stronger memory formation and in turn promote the development of pathological craving and problems in control/time management (Cox & Witten, 2019; Dolan & Dayan, 2013).

In contrast to our expectations, we did not find associations between levels of IUD and neural activity during affective go/no-go or pain empathy. Previous studies reported inhibitory and cognitive control deficits in the context of deficient frontal engagement in IUD, in particular IGD (e.g. Kuss, Pontes, & Griffiths, 2018). These findings were primarily based on samples with manifest IUD, while our findings were obtained in subjects with comparably lower symptom load suggesting that frontal deficits during inhibitory control may either represent a predisposition for rather than a consequence of excessive Internet use or alternatively may only manifest at later stages of the disorder. In line with accumulating evidence from prospective longitudinal intervention studies reporting no effects of gaming on emotional or pain empathic processes (Gao et al., 2017; Kühn et al., 2018; Yu et al., 2021), we did not observe alterations in the pain empathic and social processing domain. This may reflect that specifically the processing of Internet-or game-related stimuli is biased in IUD. In line with this explanation previous studies observed altered processing of internet related stimuli but not of general emotional or monetary reward related processing (Starcke et al., 2018; Yao et al., 2020; Yu et al., 2021).

The present findings underscore the importance of considering symptom-domains separately rather than the total symptom load of IUD related scales. While the two-factorial structure of the IAT (Pawlikowski et al., 2013) and separate diagnostic symptom domains have been emphasized, the vast majority of IUD or IGD studies determined samples based on the total symptom score. The total score may have a high clinical utility to identify individuals in need for intervention (Pontes & Griffiths, 2015; Pontes, Kiraly, Demetrovics, & Griffiths, 2014), however, the present results underscore that the identification of the precise neurocognitive and brain-based mechanisms will require a symptom-domain specific approach because distinct alterations may underlie specific behavioral dysregulations in IUD.

Findings of the present study need to be considered within the context of limitations. Firstly, IUD was associated with inhibitory control performance, but a comprehensive neurocognitive characterization in association with the IAT was not concluded. Secondly, we cannot exclude that the s-IAT sub-facets would exhibit associations with neural activity during other cognitive and emotional processes. The symptom-subdomain specific alterations need to be established across different IUD subforms such as IUD.

## Conclusions

Our findings provided the first evidence for symptom-domain specific associations with progressive alterations in the intrinsic structural and functional organization of the brain, in particular the striatal systems involved in reward, habitual and cognitive control processes. Our findings suggest that different symptom domains in IUD are neurally underpinned by separable alterations.

## Acknowledgements and funding

This work was supported by the National Key Research and Development Program of China (grant number: 2018YFA0701400). The authors declare no competing interests.

## Notes

**Funding sources** The study was supported by the National Key Research and Development Program of China (grant number: 2018YFA0701400).

**Conflict of interest** The authors declare no conflict of interest

### Competing Interest Statement

The authors have declared no competing interest.

