## supplements for "Opposing associations of Internet Use Disorder symptom domains with structural and functional organization of the striatum: a dimensional neuroimaging approach"

**Supplementary materials**

**Yu et al.,**
**Contact**

**Participants**

A comparably large sample of N = 250 subjects underwent assessment of levels of Internet Addiction Disorder via the s-IAT and multimodal MRI acquisition, including task paradigms, resting state and brain structural assessments. Aligning the questionnaire and MRI data yielded a sample of N = 219 individuals with complete data. Based on further quality assessments data from n = 2 participants were excluded due to technical failure during resting state data acquisition and data from n = 3 was excluded due to excessive head motion during resting state MRI acquisition (>3.0 mm and 3.0 degree). Based on quality assessments of the brain structural data from n = 11 subjects were additionally excluded due to a mean correlation below 2 standard deviations (SD) with respect to homogeneity of structural data reported by the CAT toolbox (http://www.neuro.uni-jena.de/cat/). A total of N = 203 subjects was included in the final multimodal data analyses (104 males; mean age ± SD = 21.65 ± 2.37 years).

**MRI acquisition protocols**

MRI data were collected using a 3 Tesla, GE Discovery MR750 system (General Electric Medical System, Milwaukee, WI). The acquisition parameters were identical for resting state and task paradigms: repetition time, 2000ms; echo time, 30ms; slices, 39; slice-thickness, 3.4mm; gap, 0.6mm; field of view, 240 × 240 $mm^2$; matrix size, 64 × 64; flip angle, 90°. For resting data, 200 volumes images were obtained for each subject. For task paradigms 218 and 488 functional volumes of T2*-weighted echo planar images were obtained for the empathy task and Go/No-go task respectively. High-resolution whole-brain T1-weighted images were acquired to improve normalization of the functional images and to determine brain structural variations (spoiled gradient echo pulse sequence with oblique acquisition, acquisition parameters: repetition time, 6ms; echo time, minimum; flip angle, 9°; field of view =256 × 256mm; acquisition matrix, 256 × 256; thickness, 1mm; 156 slices). OptoActive MRI headphones (http://www.optoacoustics.com/) were used to reduce acoustic noise exposure during fMRI.

**Resting state data acquisition**

The task paradigms were preceded by a resting state fMRI acquisition during which subjects were instructed to think of nothing in particular while keeping their eyes open and to not fall asleep. The acquisition duration was approx. 7 min.

**Pain empathy task paradigm**

Pain empathy related neural activity was assessed by means of a modified version of a previously validated blocked design pain empathy paradigm (e.g. Xu et al., 2020; Yao et al., 2016). Participants were required to view two types of pain-associated stimuli (physical pain empathy stimuli, depicting noxious stimulation of body limbs, affective pain empathy stimuli, depiction painful facial expressions) as well as corresponding non-painful control stimuli (see main text figure 1. Physical pain pictures depicted a person's hand or foot in a painful everyday situations form a first-person perspective (e.g. cutting a hand with a knife, the matched non-painful control pictures depicted e.g. cutting vegetables with a knife). The affective stimuli consisted of painful neutral facial expressions displayed by 16 Chinese subjects (8 males) (Li et al., 2019; Yao et al., 2016). Per condition 16 stimuli were included (16 per experimental condition: physical pain/affective pain/physical control/affective control) leading to a total of 64 stimuli presented over 16 blocks (4 blocks per condition) with 4 condition-specific stimuli per block (each presented for 3s). The blocks were interspersed by a jittered inter-block (8-12s) and the total duration of the paradigm was 436s acquired in a single fMRI run (further details see also Li et al., 2019).

**Affective Go/No-go task paradigm**

A modified version of a previously validated mixed event-related block design emotional (linguistic) Go/No-go paradigm (Goldstein et al., 2007; Liu et al., 2021; Protopopescu et al., 2005; Zhuang et al., 2021) was employed. During the fMRI acquisition participants were presented a sequence of words and were required to (silently) read the words and to respond as quickly and accurately depending on the font of the word stimuli. Stimuli presented in normal front were designated as Go trials and participants were required to respond via right index finger button-press while stimuli presented in italicized font were designated as No Go trials and subjects were required to withhold their response towards these stimuli. Emotional context was modulated by including blocks during which neutral, positive and negative Chinese words (matched for length, length = four characters, word frequency) presented. The paradigm encompassed 24 blocks that were presented in 2 runs with 12 blocks each. Each stimulus was presented for 300ms, followed by an inter-stimulus-interval (ITI) of 900ms, leading to a block duration of 21.6s. During each go block, 18 words in normal font (100% Go trials) were presented while during no-go blocks 12 normal font words (66.7% Go trials) and 6 italicized font words (33.3% No Go trials) were presented. Total duration of the paradigm was 16 minutes. Details see also Zhuang et al., 2021.

**Resting data preprocessing**

Resting functional MRI data were processed using SPM12b software (Wellcome Trust Center of Neuroimaging, University College London, London, United Kingdom) and DPABI software (Yan et al., 2016, http://rfmri.org/dpabi). The first 10 volumes were excluded to allow magnet steady images and active noise cancelling. The subsequent preprocessing encompassed a standard pipeline including slice timing, realignment, co-registration to the T1-weighted structural images, normalized to Montreal Neurological Institute (MNI) standard space, normalization at 3mm voxel size resolution, nuisance covariates regression (white matter, CSF, global signal), and smoothing with 8mm FWHM, band-pass filter (0.01-0.1Hz). For the connectivity analysis n = 3 subjects were excluded due to excessive head motion (exclusion criterion> 3 mm and/or 3 degrees).

**Task fMRI data preprocessing**

For task fMRI data preprocessing, the first ten volumes were discarded. The remaining functional images were subjected to a standard preprocessing pipeline including realignment to correct for head motion, co-registration to the T1-weighted structural images, normalized using the segmentation parameters from the structural images to Montreal Neurological Institute (MNI) standard space, resampling at $3mm^3$ voxel size and spatially smoothing using a Gaussian kernel with full-width at half-maximum (FWHM) of 8mm.

**Internet Addiction Test (s-IAT) distribution and psychometric properties**

Testing the normal distribution of the questionnaire data by means of Shapiro-Wilk test indicated that the distribution of s-IAT total scores and the two subscale scores were not normally distribution (details see table S2), visual inspection of the distribution (see figure S1) and examination of skewness and kurtosis (Shapiro-Wilk's W within -1 to 1, (Howell, 2012) revealed that the data displayed no strong violation of the normal distribution thus parametric tests were employed to examine associations between s-IAT scores and neural and neurocognitive indices. Psychometric properties in terms of reliability for the s-IAT were high such that the Cronbach's α of the total s-IAT was 0.86, and the Cronbach's α of loss of control/time management subscale was 0.786, the Cronbach's α of craving/social problem subscale was 0.761.

### Supplementary figures and tables

Figure S1 Distribution of IAT total scores and two subscale scores

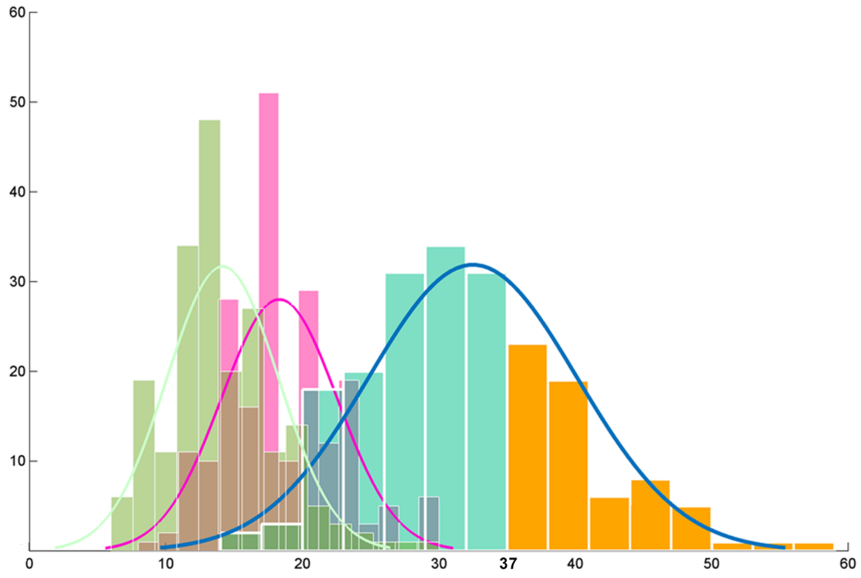


In present sample, testing the normal distribution of the questionnaire data by means of Shapiro-Wilk test indicated that the distribution of s-IAT total scores and the two subscale scores were not normally distributed. However, examination of skewness and kurtosis (Shapiro-Wilk's W within -1 to 1) revealed that the data displayed no strong violation of the normal distribution (more details see supplementary materials and table S2). Around 25% percent of whole sample (43 participants) can be defined as pathological Internet users due to s-IAT total scores exceed 37 (75 percentile was 37, blue part of IAT total scores).

Cyan represents the whole sample (N = 203), orange represents the participants (N = 48) who fulfill the criteria for pathological Internet use. Green represents the distribution of craving/social problem subscale and magenta represents the distribution of loss of control/time management subscale.

Figure S2 Activations in two tasks.

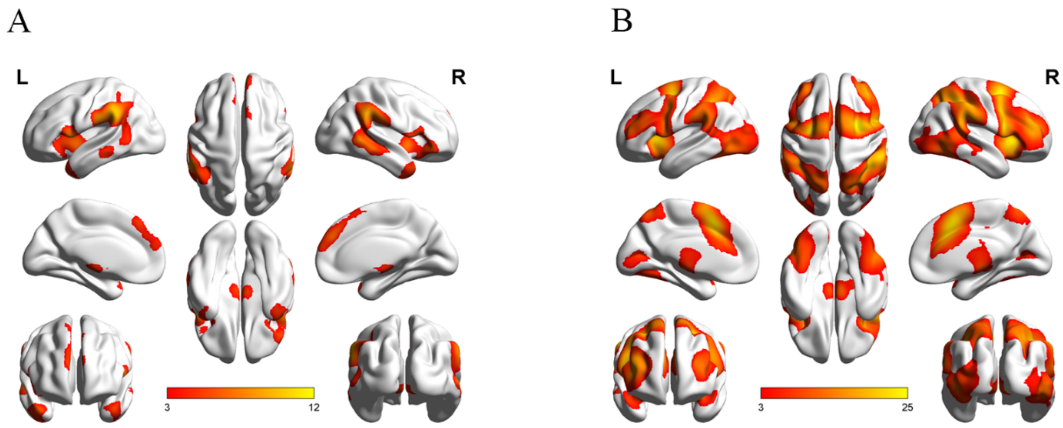


Activation in two tasks. A: The pain empathy task engaged the typical networks located in the temporal lobe, parietal lobe and frontal lobe, including bilateral superior temporal gyrus and middle temporal gyrus, bilateral inferior parietal lobe, bilateral inferior frontal gyrus and insula, right medial frontal gyrus and superior frontal gyrus (FDR $p$ <0.05, cluster size > 297 voxels). B: Activations of Go/No go task indicated that inhibitory control revealed higher activity during inhibition in a bilateral frontoparietal control network, bilateral visual network, bilateral medial prefrontal cortex, bilateral anterior cingulate cortex (FDR $p$ < 0.05).

Table S1 Short Internet Addiction Test (s-IAT) in English and Chinses language

| | English | Chinese |
|---|---|---|
| 1. | How often do you find that you stay online longer than you intended? | 你经常会发现自己在网上呆的时间比你预期的要长吗？ |
| 2. | How often do you neglect household chores to spend more time online? | 你经常会为了在网上多呆一会儿而忽略日常事务吗？ |
| 3. | How often do your grades or school work suffer because of the amount of time you spend online? | 你经常会因为上网用了很多时间而耽误了成绩或者大学的功课吗？ |
| 4. | How often do you become defensive or secretive when anyone asks you what you do online? | 你经常会在别人问起你在网上做什么的时候变得防御或者遮遮掩掩吗？ |
| 5. | How often do you snap, yell, or act annoyed if someone bothers you while you are online? | 你经常会因为别人打扰你上网而呵斥、大叫或者作出生气的行为吗？ |
| 6. | How often do you lose sleep due to being online late at night? | 你经常会因为上网到半夜而失眠吗？ |
| 7. | How often do you feel preoccupied with the Internet when offline, or fantasize about being online? | 你经常会在线下的时候总想着上网的事情或者想象着在网络上吗？ |
| 8. | How often do you find yourself saying “just a few more minutes” when online? | 你经常会发现在上网时你对自己说“再多玩几分钟”吗？ |
| 9. | How often do you try to cut down the amount of time you spend online and fail? | 你经常试图减少上网的时间但是却以失败告终吗？ |
| 10. | How often do you try to hide how long you’ve been online? | 你经常会尝试隐瞒实际在网上呆的时间吗？ |
| 11. | How often do you choose to spend more time online over going out with others? | 你经常会在网络上花费的时间比与他人一起外出的时间更多吗？ |
| 12. | How often do you feel depressed, moody, or nervous when you are offline, which goes away once you are back online? | 你经常会因为不能上网而感到郁闷、不高兴或焦虑，而一旦能够玩游戏时这些感受就烟消云散了吗？ |

*Note*: Participants must answer all items on a 5-point Likert scale: 1 = never/从不, 2 =

rarely/很少, 3 = sometimes/有时, 4 = often/经常, 5 = very often/非常常见. The total score is sum of 12 items; Loss of control/time management sub-scale includes items: 1, 2, 3, 6, 8, 9; Craving/social problems sub-scale includes items: 4, 5, 7, 10, 11, 12.

Table S2 distribution description of questionnaire data

| | s-IAT total score | Loss of control subscale score | Craving/social problem subscale score |
|---|---|---|---|
| N | | 203 | |
| Mean (SD) | 32.48 (7.63) | 18.32 (4.24) | 14.17 (4.09) |
| total | 60 | 30 | 30 |
| Median | 32 | 18 | 14 |
| Minimum | 14 | 8 | 6 |
| Maximum | 59 | 30 | 30 |
| $75^{th}$ percentile | 37 | 21 | 16 |
| Skewness (SE) | 0.57 (0.17) | 0.37 (0.17) | 0.69 (0.17) |
| Kurtosis (SE) | 0.49 (0.34) | -0.02 (0.34) | 1.02 (0.34) |
| Shapiro-Wilk W | 0.98 | 0.98 | 0.96 |
| Shapiro-Wilk *p* | 0.005 | 0.008 | < 0.001 |

Table S3 the partial correlation between questionnaire data and Go/no-go behavioral data.

| | condition | | s-IAT total score | Loss of control subscale score | Craving/social problem subscale score |
|---|---|---|---|---|---|
| Accuracy | Negative go | Pearson's *r* | -0.131 | -0.105 | -0.136 |
| | | *p* | 0.06395 | 0.13723 | 0.05380 |
| | Positive go | Pearson's *r* | -0.061 | -0.054 | -0.059 |
| | | *p* | 0.38720 | 0.44929 | 0.40261 |
| | Neutral go | Pearson's *r* | -0.033 | -0.048 | -0.013 |
| | | *p* | 0.64136 | 0.50189 | 0.85424 |
| | Negative no-go | Pearson's *r* | -0.215* | -0.186* | -0.211* |
| | | *p* | 0.00215 | 0.00823 | 0.00266 |
| | Positive no-go | Pearson's *r* | -0.233** | -0.189* | -0.241** |
| | | *p* | 0.00087 | 0.00717 | 0.00058 |
| | Neutral no-go | Pearson's *r* | -0.203* | -0.162 | -0.211* |
| | | *p* | 0.00391 | 0.02137 | 0.00258 |
| Reaction time | Negative go | Pearson's *r* | -0.157 | -0.131 | -0.158 |
| | | *p* | 0.02629 | 0.06363 | 0.02520 |
| | Positive go | Pearson's *r* | -0.139 | -0.120 | -0.137 |
| | | *p* | 0.04832 | 0.09062 | 0.05272 |
| | Neutral go | Pearson's *r* | -0.130 | -0.107 | -0.134 |
| | | *p* | 0.06504 | 0.13138 | 0.05856 |

*Note:* We calculated the partial correlation between the scores and accuracy and reaction time of go and no-go trials for three conditions (negative, positive, neutral) separately and used age and gender as covariates. *Note:* used Bonferroni correction for multiple comparison, * $p < 0.0083$ (0.05/6), ** $p < 0.0016$ (0.01/6), *** $p < 0.00016$ (0.001/6).

Table S4 Subregions in Brainnetome Atlas that served as seed regions-of-interest (ROIs)

| Seed ROIs | subregions | Label ID | MNI (x,y,z) |
|---|---|---|---|
| Ventral striatum (VS) | left ventral caudate | 219 | -12, 14, 0 |
| | right ventral caudate | 220 | 15, 14, -2 |
| | left nucleus accumbens | 223 | -17 ,3, -9 |
| | right nucleus accumbens | 224 | 15, 18, -9 |
| Dorsal striatum (DS) | left ventral medial putamen | 225 | -23, 7, -4 |
| | right ventral medial putamen | 226 | 22, 8, -1 |
| | left dorsal caudate | 227 | -14, 2, 16 |
| | right dorsal caudate | 228 | 14, 5, 14 |
| | left dorsolateral putamen | 229 | -28, -5, 2 |
| | right dorsolateral putamen | 230 | 29, -3, 1 |

Table S5 Activation during the empathy task

| Task | MNI coordinate of peak intensity | Number of voxels | Hemisphere | S |
|---|---|---|---|---|
| Pain stimuli > Non-pain stimuli | 63, -39, 33 | 3129 | right | Middle Temporal Gyrus (MTG), Superior Temporal Gyrus (STG), Inferior Parietal Lobe (IPL), Inferior Frontal Gyrus (IFG), inferior orbital frontal gyrus, Insula |
| | -63, -39, 33 | 1504 | left | IPL |
| | -51, 6, 3 | 1187 | left | STG, IFG, Insula |
| | 9, 45, 30 | 651 | right | Medial Frontal Gyrus (MFG) |

*Note:* use FDR $p < 0.05$ for multiple comparison. Cluster size > 297 in pain empathy.

Table S6 Regions of overlap between caudate and bilateral ventral and dorsal striatum

| Cluster A | Cluster B | Overlap region | MNI coordinate | Number of voxels |
|---|---|---|---|---|
| Caudate | Bilateral ventral striatum | right caudate | 7.5, 7.5, -1.5 | 62 |
| | | left caudate | -4.5, 6, -1.5 | 91 |
| | Bilateral dorsal striatum | right caudate | 9, 6, 3 | 288 |
| | | left caudate | -9, 3, 6 | 124 |
